# Systems approaches provide new insights into *Arabidopsis thaliana* root growth under mineral nutrient limitation

**DOI:** 10.1101/460360

**Authors:** Nadia Bouain, Arthur Korte, Santosh B. Satbhai, Seung Y. Rhee, Wolfgang Busch, Hatem Rouached

## Abstract

The molecular genetic mechanisms by which plants modulate their root growth rate (RGR) in response to nutrient deficiency are largely unknown. Using a panel of *Arabidopsis thaliana* natural accessions, we provide a comprehensive combinatorial analysis of RGR variation under macro- and micronutrient deficiency, namely phosphorus (P), iron (Fe), and zinc (Zn), which affect root growth in opposite directions. We found that while -P stimulates early RGR of most accessions, -Fe or -Zn reduces it. The combination of either -P-Fe or -P-Zn leads to suppression of the growth inhibition exerted by -Fe or -Zn alone. Surprisingly, Arabidopsis reference accession Columbia (Col-0) is not representative of the species under -P and -Zn. Using a genome wide association study, we identify candidate genes that control RGR under the assayed nutrient deficiency conditions. By using a network biology driven search using these candidate genes, we further identify a functional module enriched in regulation of cell cycle, DNA replication and chromatin modification that possibly underlies the suppression of root growth reduction in -P-Fe conditions. Collectively, our findings provide a framework for understanding the regulation of RGR under nutrient deficiency, and open new routes for the identification of both large effect genes and favorable allelic variations to improve root growth.

## Introduction

Global climate change and population increase pose a tremendous challenge and prompt an urgent need for efficient agriculture and food production. The current projection is that the world population will be over 9.8 billion by 2050, and global food production will have to further increase by 70% to sustain this population (Tomlinson, 2013). At the same time, climate change is associated with a decrease of important micronutrients such as iron (Fe) and zinc (Zn) in staple foods such as rice (Zhu et al., 2018). The bio-availability of Fe and Zn is often limited in the soil, leading to reductions in growth, crop yield, and quality. As plants are the entry point of these elements in the food web, low accumulation of these elements in plants is associated with malnutrition in humans. Zn and Fe deficiencies are estimated to affect up to 2 billion people worldwide (Hilty et al., 2010). Moreover, modern industrialized agriculture has created a strong demand and dependency for fertilizers, posing the danger of acute limitation of non-renewable fertilizer components, in particular phosphorus (P) (Abelson 1999; Cordell et al., 2009; Neset and Cordell 2011). All these nutrients (P, Zn and Fe) are taken-up by plants at the root-soil interface, and plants often face conditions in which one or more of these elements are limiting (Shahzad et al., 2014; Bouain et al., 2016). Thus, improving the capacity of plants to absorb these nutrients from the soil is a major goal of crop improvement. The development of crops with higher tolerance for individual and multiple nutrient stresses is a direction towards more sustainable and efficient agriculture.

P is an essential macronutrient for plant growth and development. P is a critical component of many macromolecules (e.g. DNA), energy sources (e.g. ATP) and regulation of signal transduction via phosphorylation (Poirier and Bucher, 2002; Rouached et al., 2010). Plants acquire P as soluble inorganic phosphate (Pi) by a suite of high affinity phosphate transporters in the root (Secco et al., 2017). In soil, P distribution is heterogeneous (Hanlon et al., 2018), and usually found in shallow soil layers (Heppell et al., 2016). To cope with P heterogeneity in soil, plants increase root growth in shallow soil layers that promote topsoil foraging, thereby conferring an advantage for P acquisition (Lynch, 2011; Miguel et al., 2013).

The effects of P deficiency (-P) on the root system have been studied in many plant species. For the model plant *Arabidopsis thaliana* (*A. thaliana*), the accession Columbia (Col-0) is commonly considered as the reference (Somssich et al., 2018). Changes that occur in root architecture in -P consist of reduction of primary root growth (PRG), an increase in the growth of secondary roots, and an increase in root hair length and density (for review, Bouain et al., 2016). A handful of genes involved in PRG under -P have been cloned. Under -P, mutants of these genes are characterized either by their ability to maintain PRG such as the *low phosphate root 1* (*lpr1*) mutants (Svistoonoff et al., 2007), or by a strong inhibitory effect of the PRG (hypersensitivity) such as the *phosphate deficiency response 2* (*pdr2*) mutants (Ticconi et al., 2004) and *hypersensitive to pi starvation 7* (*hps7*) (Kang et al., 2014). More genes involved in inhibition of PRG were recently identified, including a transcription factor SENSITIVE TO PROTON RHIZOTOXICITY (STOP1) and its target ALUMINUM ACTIVATED MALATE TRANSPORTER 1 (ALMT1) (Mora-Macías et al., 2017; Balzergue et al., 2017). In addition to genetic screens for mutants affected in their response to -P, *A. thaliana* has natural accessions that are either oversensitive (e.g. Shadara) or more tolerant (e.g. Landsberg *erecta*) to -P compared to Col-0 (Reymond et al., 2006). The existence of natural variation in root growth in response to -P indicates the potential for discovering new genes that control this trait via quantitative genetic approaches.

Fe and Zn are involved in vital biological processes ensuring proper functioning of the cell. Fe is a cofactor for numerous enzymes and a part of Fe–S clusters that are a major sink for Fe and essential for many important cellular processes such as photosynthesis and respiration (White and Broadley, 2009; Couturier et al., 2013). Similarly, Zn functions as a cofactor for hundreds of enzymes (Sinclair and Krämer, 2012). Since Zn and Fe are taken up by roots, improvement of root growth could help increase Zn and Fe content in plants. In Col-0, deficiencies in Fe and Zn impose a change in root architecture in a contrasting manner (Gruber et al., 2013; Bouain et al., 2018). While Fe deficiency (-Fe) inhibits root elongation (Gruber et al., 2013; Satbhai et al. 2017), Zn deficiency (-Zn) slightly promotes early root growth (Bouain et al., 2018). Only few genes that regulate primary root growth under -Fe or -Zn conditions have been identified. For instance, mutations in the BASIC HELIX-LOOP-HELIX-TYPE transcription factors POPEYE (Long et al., 2010), or the two interacting transcription factors, bHLH34 and bHLH104 (Li et al., 2016), inhibited the primary root growth in -Fe. More genes that control root architecture under -Fe and -Zn conditions remain to be identified. The convenience of genome-wide association studies (GWAS) in *A. thaliana* offers an opportunity to explore the diversity in the response of accessions to mineral nutrients and interactions between nutrients, which promises the identification of candidate genes and their genetic variants controlling these traits.

Recent research has shown that nutrient homeostasis by interaction between nutrients is a general rule in plants rather than an exception (for review, Briat et al., 2015; Rouached and Rhee, 2017). P and Zn or P and Fe interact in the plant, and these interactions are visible at the molecular level as revealed by transcriptome analyses, where the deficiency of one-element induces or represses genes involved in the regulation of the other element (Misson et al., 2004; Kellermeier et al., 2014; Bouain et al., 2014; Briat et al., 2015; Li and Lan, 2015). The response to -P or -Fe may share common hormone signals, such as cytokinin. For example, cytokinin signaling, mediated by the cytokinin receptors CYTOKININ RESPONSE 1/WOODEN LEG/ARABIDOPSIS HISTIDINE KINASE 4 (CRE1/WOL/AHK4), is involved in the response to -P or -Fe in *A. thaliana* (Col-0) (Martin et al., 2000;Franco-Zorrilla et al., 2005; Séguéla et al., 2008). The outcome of the interaction of these nutrients is also visible at the morphological level (Kellermeier et al., 2014). Perhaps the most prominent example is the -P-Fe interaction and its effect on root growth. Root growth inhibition in response to -P has been proposed to be the result of Fe “toxicity” (Ward et al., 2008; Bournier et al., 2013) and it has been suggested that the inhibition of Arabidopsis (Col-0) primary root growth under -P is due to a presumed overabundance of available Fe, and is not solely due to –P alone (Ward et al., 2008; Bournier et al., 2013). This Fe overabundance was proposed to depend on malate exudation and presumably malate chelating Fe. The malate transporter ALMT1 was shown to be involved in this process, likely by promoting Fe accumulation in the root meristem causing the inhibition of cell expansion under -P (Balzergue et al., 2017; Mora-Macias et al., 2017). Fe accumulation in the root tip of P-deficient plants was proposed to be the cause for the differentiation of the apical root meristem - possibly through a prevention of the symplastic cell-to-cell communication as a consequence of callose deposition (Mora-Macias et al., 2017). The PDR2–LPR1 module was proposed to mediate this callose accumulation in root meristems experiencing -P (Balzergue et al., 2017). It has been proposed that the CLAVATA3/ESR (CLE)-related protein 14 precursor (CLE14) is the signal that triggers full root meristem differentiation in -P through CLV2/PEPR2 receptors (Gutiérrez-Alanís et al., 2017). It has been shown that the CLE14 pathway acts downstream of LPR1/LPR2 (Gutiérrez-Alanís et al., 2017). In contrast, these cellular hallmarks of -P were not observed under simultaneous P and Fe deficiency. The callose deposition was not detected and root growth was comparable to plants grown under complete medium. Thus far, these two modules LPR1-PDR2 and STOP1-ALMT1 have been used to explain how P and Fe signals shape root growth under P and Fe deficient conditions (Balzergue et al., 2017; Mora-Macias et al., 2017; Abel et al., 2017; Gutierrez-Alanis et al., 2018).

Plants have evolved mechanisms to sense and respond to nutrient deficiency early in their life cycle. Soon after germination, roots are the main targets of nutrient deficiency stress. Root growth responses to nutrient changes are genetically determined, and vary between and within plant species (Ristova and Busch, 2014). Identifying genes and mechanisms that underlie natural variation of root growth has become possible through large-scale phenotyping and various mapping approaches (Slovak et al., 2016). Here we set out to investigate variation in the root growth rate (RGR) of a panel of natural accessions in *A. thaliana* under single and combined deficiencies of P, Fe, and Zn. We used GWAS to identify candidate genes involved in RGR regulation under each of the growth conditions tested. Finally, we used a network biology driven search using these candidate genes to identify potential networks and processes that are relevant for determining RGR under the nutrient deficiency conditions. Taken together, our findings shed light on the regulation of root growth by combinatorial mineral nutrient cues and provide a foundation for guiding new agronomical and biotechnological strategies to improve root growth.

## Results

### RGR responds distinctly to single and combined nutrient deficiencies in a genotype dependent manner

Natural variation of root system architecture was reported for -P (Chevalier et al., 2003; Reymond et al., 2006; Kawa et al., 2016) as well as for -Fe (Satbhai et al., 2016) and -Zn (Bouain et al., 2018), yet never for their combination (-P-Fe or -P-Zn).In particular, root growth rate (RGR) as a trait has not been evaluated under any nutritional stresses. Therefore, we set out to explore the natural variation of RGR in *A. thaliana* by investigating the RGR of 227 genetically diverse natural accessions from the RegMap population (Horton et al., 2012) (Supplemental Table 1) grown on different media. We tested six growth conditions: control (Ct), deficiency of P (-P), Fe (-Fe), Zn (-Zn), P and Fe (-P-Fe), and P and Zn (-P-Zn) (Figure 1). Seedlings were imaged every day at the same time, and the primary root length (PRL) was determined using the BRAT software (Slovak et al., 2014). For each accession, we recorded the primary root length (PRL) of three, four and five-day old seedlings (Supplemental Table 1). We first examined the PRL of Columbia-0 (Col-0) accession under all growth conditions tested, as this accession is most widely used in Arabidopsis research. Consistent with previous studies (Gruber et al., 2013; Satbhai et al. 2017; Bouain et al., 2018), -P or -Fe caused a reduction of the Col-0 PRL, while -Zn slightly promoted it compared to the Ct condition (Figure 2). Also consistent with previously published results (Ward et al., 2008), the reduced PRL observed in -P was not observed in -P-Fe. A similar suppression of -P dependent growth rate reduction was observed in -P-Zn (Figure 2). Overall, the response of PRL in Col-0 was consistent with previous reports, thus validating our experimental setup.

**Figure 1.**
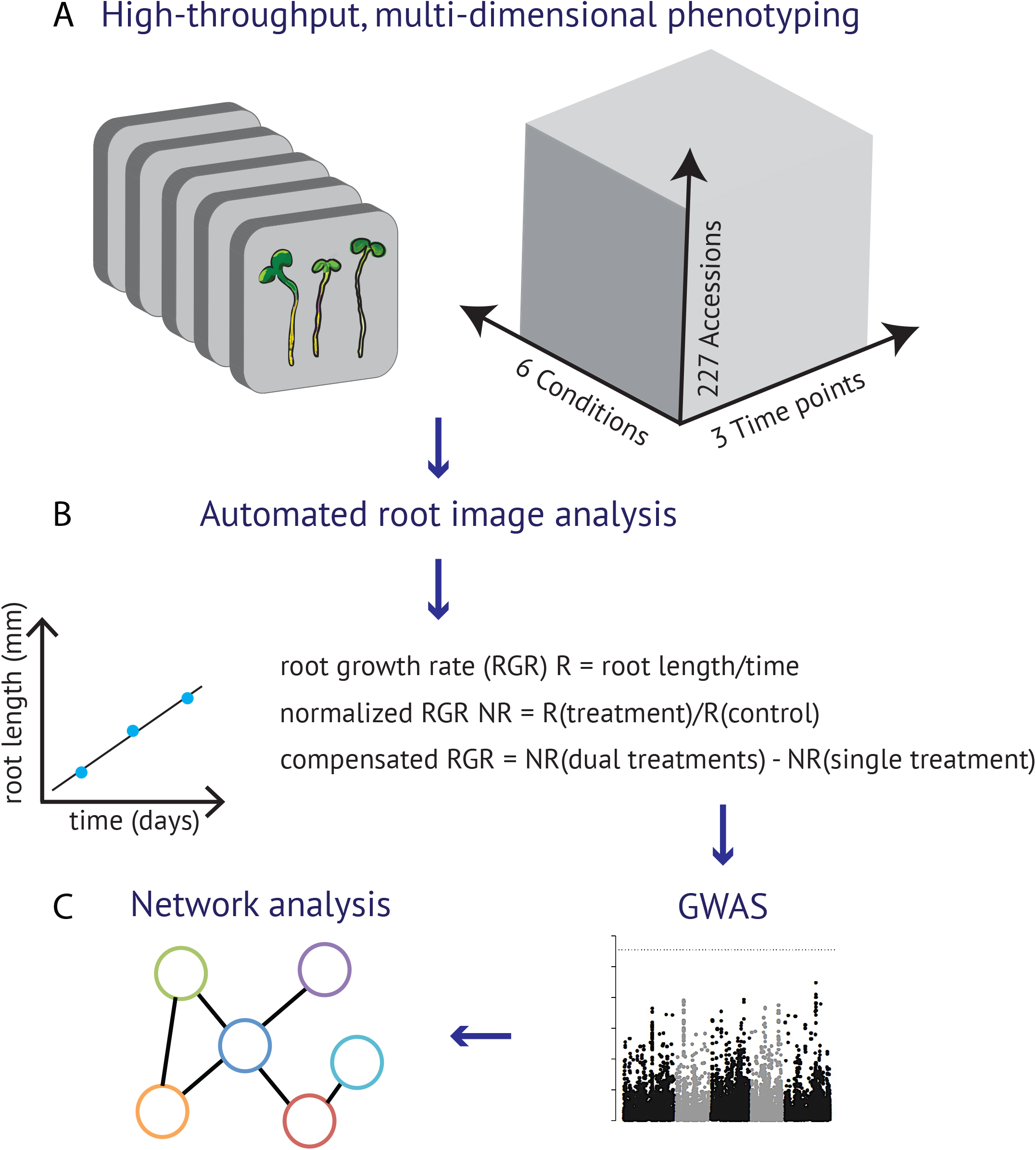
A systems framework to study root growth under mineral limitation. (A) Design of the experimental set up. 227 *Arabidopsis thaliana* accessions were grown on vertical agar plates containing different media, including control (Ct), deficiency of phosphorus (-P), iron (-Fe), zinc (-Zn), phosphorus and iron (-P-Fe), or phosphorus and zinc (-P-Zn). Seedlings were daily imaged and primary root length (PRL) of 3-, 4-, 5-day-old seedlings was determined. (B) The root growth rate (RGR) was determined by conducting linear regression on PRL of twelve replicates per accession and for each treatment. Five GWAS were performed using on normalized RGRs, which were obtained for each accession by calculating the ratio between RGR on each nutrient deficiency condition divided by the value on the control condition (Ct). GWAS was also performed on the variation of normalized RGR on -P-Fe and -Fe, or -P-Zn and –Zn by the value on control condition (Ct) and expressed as ∆RGR_(-P-Fe, -Fe)_ = RGR_(-P-Fe/Ct)_ - RGR_(-Fe/Ct)_, and for -P-Zn and –Zn ∆RGR_(-P-Zn, -Zn)_ = RGR_(-P-Zn/Ct)_ - RGR_(-Zn/Ct)_. (C) The significance of associations between phenotypes and the single nucleotide polymorphisms (SNPs) markers was evaluated using linear mixed model. The GWAS candidate genes for a given RGR trait were used to identify a gene connected with one another, which form a molecular pathway, using a machine learning algorithms to infer co-functional links from genomics data, AraNet (Lee et al., 2015).

**Figure 2.**
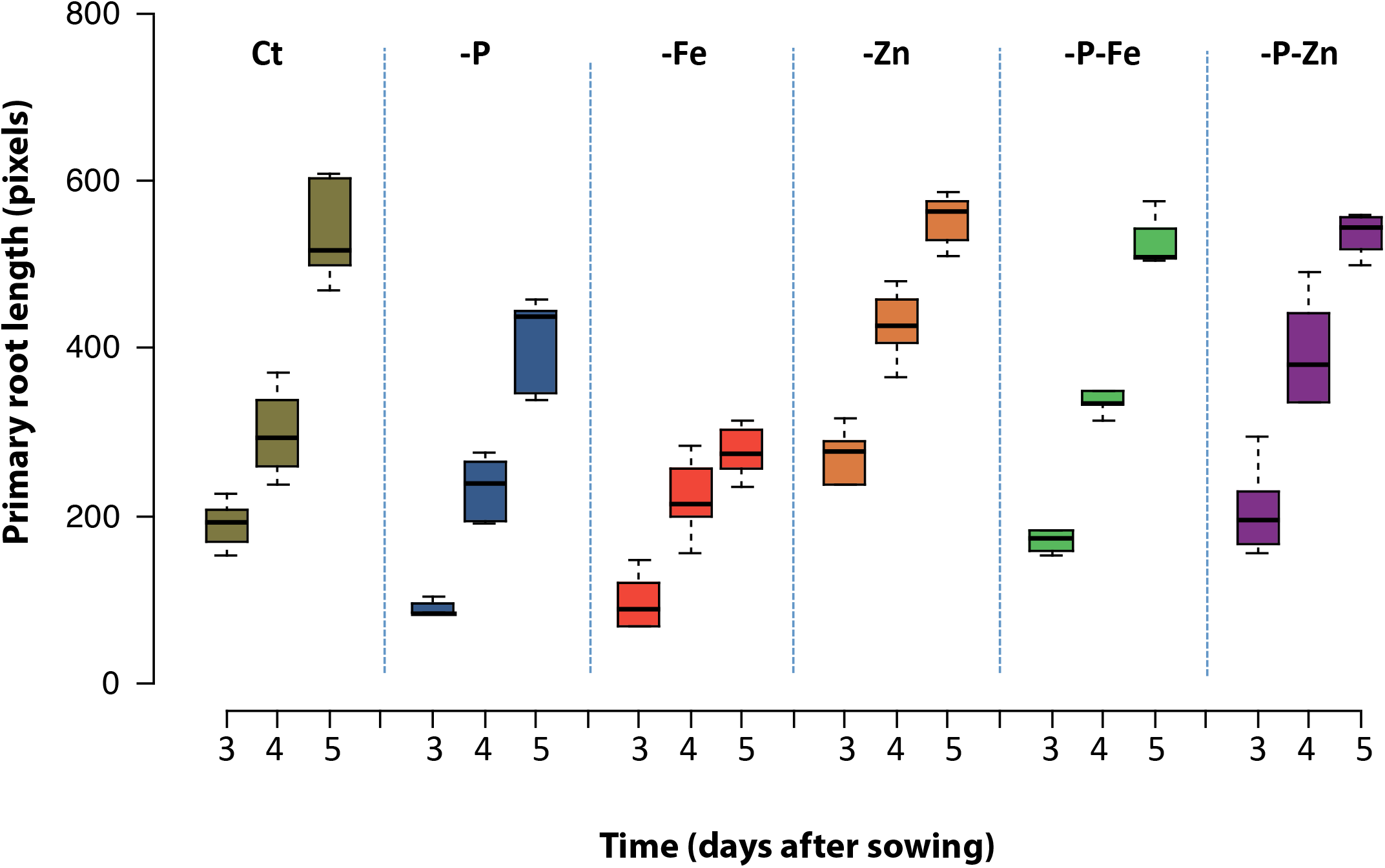
Effect of single and double deficiencies of iron, zinc and phosphorus on the primary root elongation of the *Arabidopsis thaliana* reference accession Col-0. Seeds of the *A. thaliana* Col-0 accession were germinated on six different nutrient conditions: control (Ct), deficiency of P (-P), Fe (-Fe), Zn (-Zn), P and Fe (-P-Fe), and P and Zn (-P-Zn). The primary root length was determined on 3-, 4-, and 5- day-old seedlings.

We next examined the whole set of accessions. To account for differences in germination of the accessions, we used RGR for all further comparisons. For this we conducted linear regression on the root length measurements, whereby the regression coefficient provided an estimate of the root growth rate in different treatments (Figure 1, Supplemental Table 1, 2). Under the assumption of linear growth during this early growth period, the regression coefficient on the replicates is an ideal measurement, and indeed the estimated slopes are significant for most accessions in most treatments (99.5% for -P, 98.7% for -PZn%, 97,8% for -PFe, 96,5% for Ct, 93% for -Fe, 92,5% for -P-Zn) (Supplemental Table 1). Because the data sets were obtained simultaneously, they constitute a unique resource for comparing the RGRs between accessions. Our analysis revealed a large variation of RGR among the accessions in the control condition (Ct) and in response to each of the nutrient deficiency conditions (Figure 3). To test whether the variation of root growth is genetically determined, we analyzed the heritability (H^2^) (Lynch et al., 1998) of RGR using estimates from the mixed model. We found that the phenotypic variation for RGR in response to the different conditions is a highly heritable trait (Supplemental Table 3). Like Col-0, the RGR of most accessions was reduced by -Fe treatment, and the extreme accessions included Zu-1, MIR-0, Rmx-A02, Edinburgh-5, and Ove-0. However, unlike Col-0, -Zn reduced RGR in most accessions, and the extreme accessions included Shadara, Sq-1, Sapporo-0, and Si-0. Here Col-0, which slightly increases RGR, was the exception to the rule. We found a similar surprise in -P. In -P, while Col-0 showed a reduction in their RGR, most accessions behaved in an opposite manner; their RGR was increased compared to Ct. When grown under combined nutrient deficiency (-P-Fe and -P-Zn), most accessions showed an RGR that was similar to the -P effect and distinct from -Fe or -Zn alone (Figure 3). P deficiency alleviated the RGR reduction mediated by Fe or Zn deficiency. Taken together, our analysis revealed that -P generally promoted early primary root growth whereas -Fe or -Zn reduced it in *A. thaliana.* In addition, removing P alleviated the root growth reduction brought on by Fe or Zn deficiency. This general response varied across natural accessions in response to single or combined nutrient deficiencies. Importantly, Col-0 was not the best representative accession for the responses in -P or -Zn conditions.

**Figure 3.**
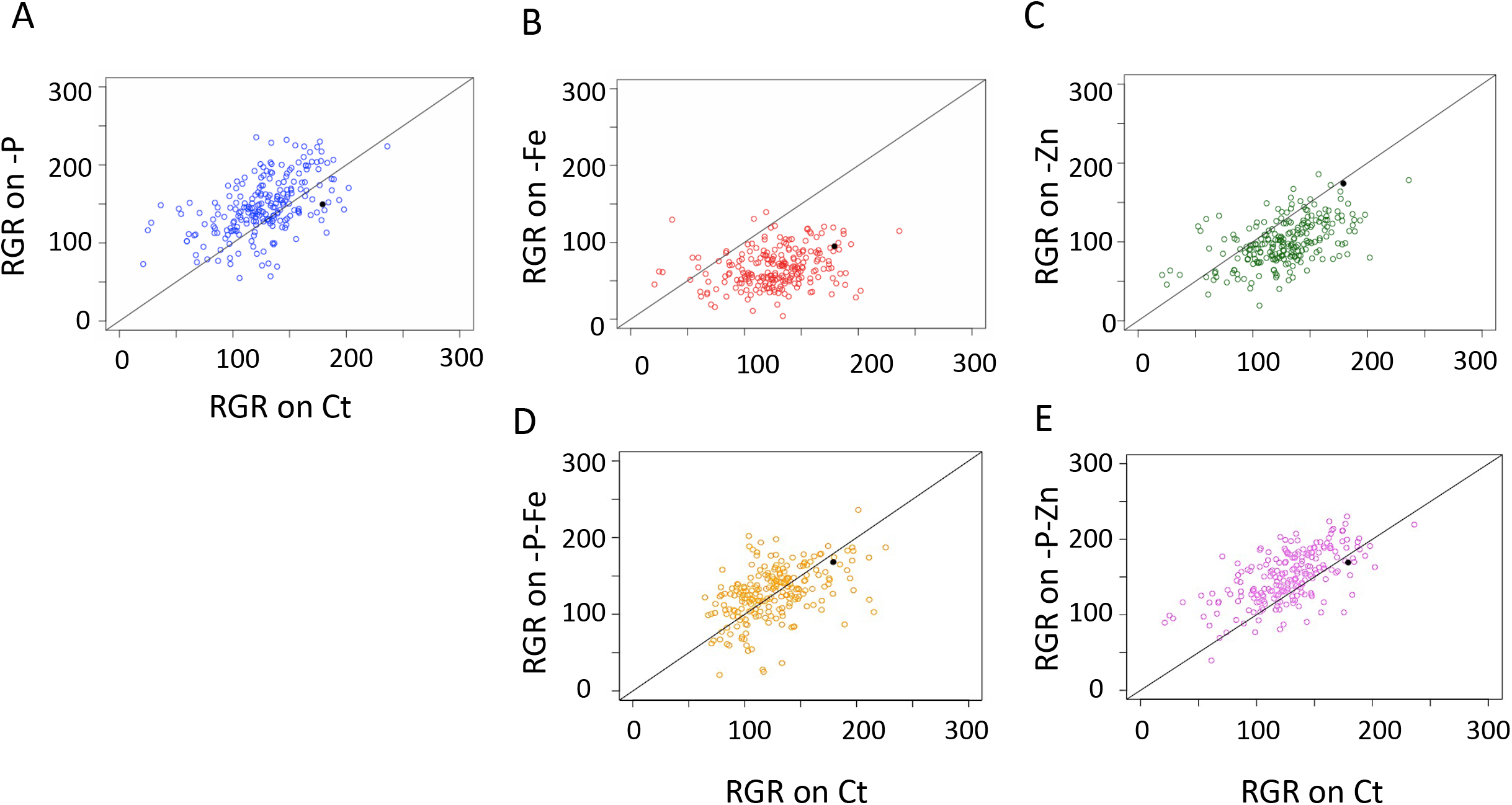
Natural variation of 227 *Arabidopsis thaliana* accessions in their responses to single and double deficiencies in iron, zinc and phosphorus. (A) Relationship between root growth rate (RGR) under phosphate deficiency (-P) and RGR under control condition (Ct). (B) Relationship between RGR under iron deficiency (-Fe) and root growth in Ct. (C) Relationship between RGR under zinc deficiency (-Zn) and RGR in Ct. (D) Relationship between RGR in -P-Fe and RGR in Ct. (D) Relationship between root growth rate under in -P-Zn and RGR in Ct. Each data point represents the RGR estimated from a pool of plants (median n in each treatment > 10). The solid black dot represents the Col-0 reference accession. The solid black line represents the diagonal (i.e. the growth that would be expected without any response) and not the results of a regression analysis.

### Distinct genetic architectures underlie the response to single and combined nutrient stresses in *Arabidopsis thaliana*

To determine the genomic loci that control RGR in response to mineral nutrient deficiencies, we performed GWAS using normalized RGRs (Figure 4, Supplemental Table 2). For each accession, normalized RGR was defined as the ratio between RGR from each nutrient deficiency condition divided by the RGR from the control condition (Ct). The significance of associations between phenotypes and the single nucleotide polymorphism (SNP) markers was evaluated using a linear mixed model (a modified version of EMMA) (Kang et al. 2008) that corrects for population structure. All considered normalized RGRs show low but a significant estimated heritability (H^2^) (Supplemental Table 3). We next used ~1.7 million markers with a minor allele frequency of at least 5% in the population and corrected the associated p-values for multiple hypothesis testing using a 5% Bonferroni threshold (Figure 4 A-E). Using this conservative threshold, and by taking into account genes present in a 10kb window surrounding the underlying significant markers - acknowledging the rapid decay of LD in the Arabidopsis population (Gan et al., 2011) - we identified a list of 145 candidate genes (Supplemental Table 4) corresponding to 87 significant SNPs in 32 distinct genomic regions (Supplemental Table 5). Of these, 96 genes (49 SNPs) were associated with only one trait, while 31 genes (19 SNPs) were associated with two traits, and 18 genes (19 SNPs) were associated with 3 traits (Supplemental Table 4). Among these candidates, some are known to be involved in regulating root growth under mineral nutrient deficiency. For instance, the *CLV2* gene is known to trigger full root meristem differentiation under -P (Gutiérrez-Alanís et al., 2017) and was identified in our GWAS RGR on -P (p-value=2.1*10^-8^). Many candidate genes related to different classes of hormone signaling were also identified, including genes involved in auxin, gibberellin, cytokinin, ABA, and brassinosteroid pathways. For instance, the gene *BRASSINAZOLE-RESISTANT 1* (*BZR1, p=1.9*10^-9^)* was identified to associate with the regulation of RGR under-Zn. Interestingly, there was a very limited overlap (if any) between the gene lists of single nutrient deficiencies (-P, -Zn, and -Fe) (Figure 4F). No common gene was detected between -P and -Fe, nor between -Fe and -Zn (Figure 4F, Supplemental Table 4). Only in the case of -P and –Zn, two overlapping regions (tagged by 17 SNPs) corresponding to 10 candidate genes were detected. The first region was significantly associated with -P-Zn, and the second region was associated with -P-Fe, in addition to -P and -Zn (Supplemental Table 6). The three associated SNPs in the first region, common between -P, -Zn, and-P-Zn, are in complete LD and tag the genes AT3G29570 and AT3G29575 (P-values: -P (p=2.1*10^-8^), -Zn (p=3.3*10^-9^), and -P-Zn (p= 9.0*10^-10^)). AT3G29575 is a member of the family of ABI five binding proteins (AFPs) and is involved in the stress response of germinating seeds and seedlings through modulation of ABA signaling (Garcia et al., 2008). Taken together, our GWAS analysis allowed the identification of interesting candidate genes that may underlie the natural variation of RGR in response to nutrient deficient conditions, and revealed that root growth responses to single stresses and to their combinations might be regulated by distinct genetic programs rather than being regulated in a simple additive manner.

**Figure 4.**
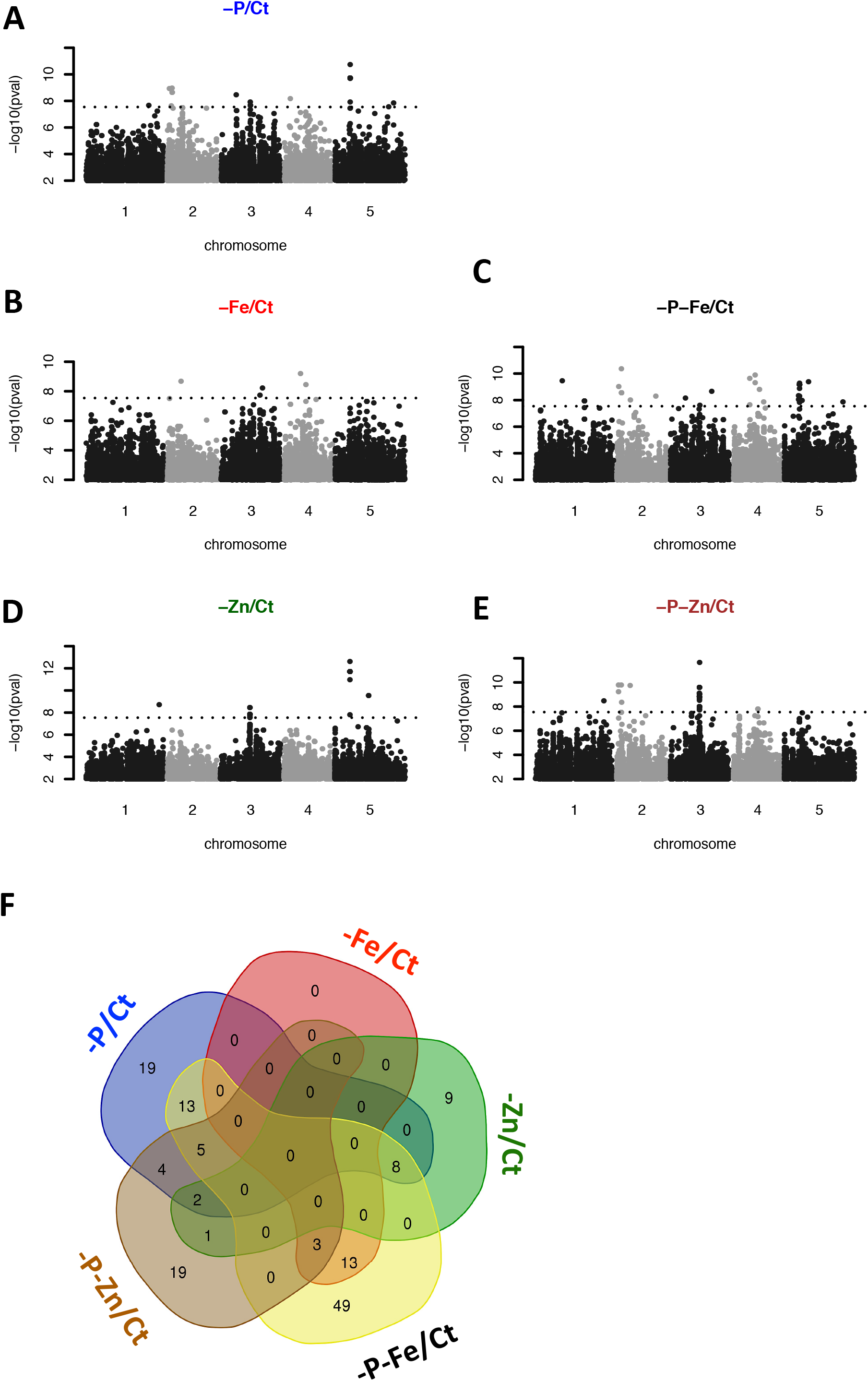
Genetic architectures of root growth rate control in response to single and double deficiencies of iron, zinc and phosphorus from GWAS. (A-E) Genome-wide distribution of the −log10 P-values of SNP/phenotype associations using a linear mixed model method that corrects for population structure (AMM). SNPs associated with the root growth rate under the deficiency of -P (A), -Fe (B) -Zn (C), -P-Fe (D), or -P-Zn (E) compared to control condition (Ct) are presented. SNPs are plotted according to their position along the chromosomes. Plotting colors alternate between black and grey in order to facilitate the visualization of each of the five chromosomes. 5% Bonferroni threshold is indicated by black dashed line. (F) Van Diagram of GWAS candidate genes underlying Arabidopsis RGR under the different nutrient-deficient conditions: -P, -Fe, -Zn, -P-Fe and -P-Zn conditions. The Van diagram was generated using a web-based tool for the analysis of sets through Venn diagrams (InteractiVenn) (Heberle et al., 2015).

### The negative regulation of RGR by iron and zinc deficiencies is suppressed when combined with phosphate deficiency

In line with the literature, we found that Col-0 plants grown under -P-Fe have longer primary roots compared to plants grown either on -P or -Fe (Figure 2). But, whether this compensation is a general mechanism of adaptation or just an exception for a few accessions, and whether -P could also alleviate -Zn’s negative effect on RGR in Arabidopsis remained unknown. We therefore set out to answer these questions. We compared normalized RGRs on combined nutrient stress (-P-Fe/Ct or -P-Zn/Ct) to normalized RGRs on single nutrient stress (-P/Ct, -Fe/Ct, or -Zn/Ct) (Figure 5 A-C). First, only a subset of accessions displayed a reduction of RGR on -P (e.g. Col-0, Sorbo, Rd-0, Rmx-A180, Mr-0, Hau-0, Got-22, Mdn-1, RRS-10) and they were compensated by the combined stress -P-Fe (Figure 5A, black dot for Col-0 and red dots for the rest). Second, reduction of RGR in -Fe was generally alleviated under combined stress of -P-Fe in *A. thaliana* accessions. Here the reference accession Col-0 behaved similarly to the majority of the accessions (Figure 5B, black dot). Finally, except for few accessions including Col-0, -P-Zn treatment generally caused a large compensation of RGR compared to -Zn (Figure 5C, black dot for Col-0). This compensation was widespread across accessions. A few examples of RGR compensation are shown for accessions such as Shadara, Sq-1, Sapporo-0, and Si-0 (Figure 6). Taken together, our results indicate that the RGR inhibition of Fe and Zn deficiencies is alleviated by P deficiency as a general mechanism in this species.

**Figure 5.**
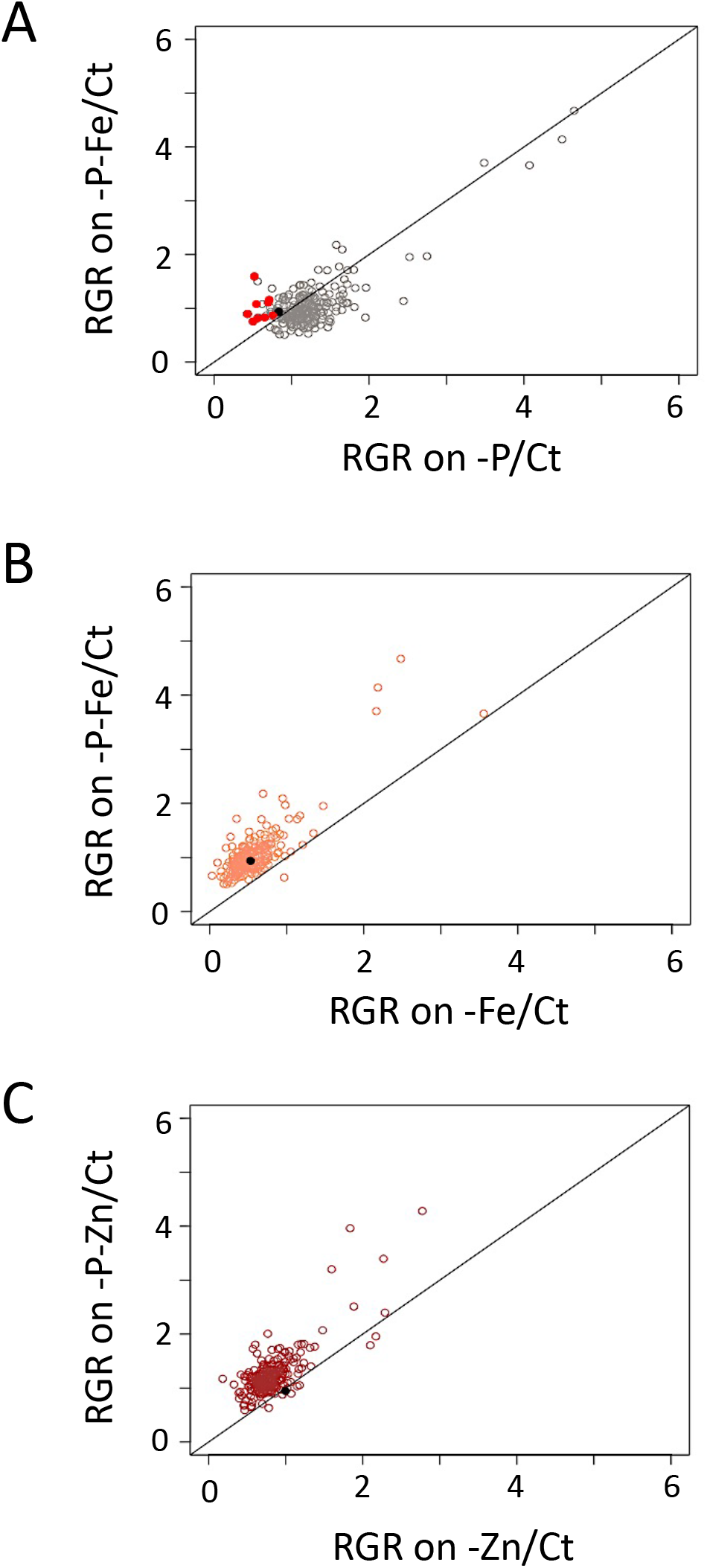
P deficiency suppresses negative effects of Fe and Zn deficiency on the root growth rate of most *Arabidopsis thaliana* accessions. The root growth rate (RGR) of 227 accessions of *Arabidopsis thaliana* grown under each of the nutrient-deficient conditions was normalized to the corresponding values under control conditions (Ct). (A) Relationship between RGR under combined phosphorus and iron deficiency (-P-Fe/Ct) and RGR under phosphorus deficiency (-P/Ct). The red dots represent the ecotypes that showed limited RGR by -P considering one standard deviation away from the mean with 15.8% confidence limit to the RGR data, and which are promoted in -P-Fe. (B) Relationship between RGR under phosphorus and iron deficiency (-P-Fe/Ct) and RGR under iron deficiency (-Fe/Ct). (C) Relationship between root RGR under phosphorus and zinc deficiency (-P-Zn/Ct) and RGR under zinc deficiency (-Zn/Ct). Each data point was obtained from the analysis of RGR from a pool of plants (n ≥ 9). The solid black dot represents the Col-0 ecotype. The solid black lines represent the diagonal.

**Figure 6.**
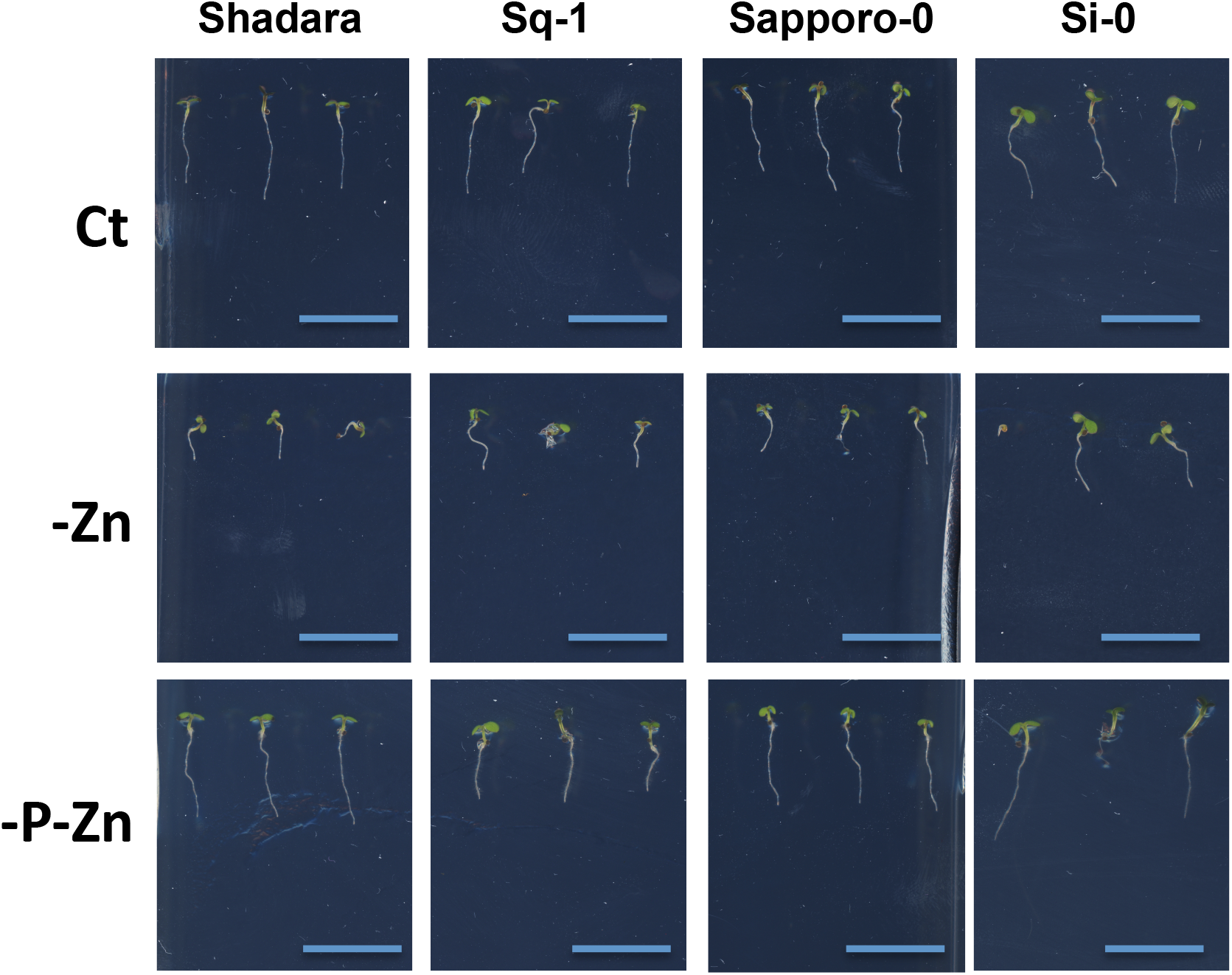
Combined phosphorus and zinc deficiency promotes root growth rate of most *Arabidopsis thaliana* accessions. Representative images of contrasting primary root growth phenotype (5 days) of four *Arabidopsis thaliana* accessions (Shadara, Sq-1, Sapporo-0, Si-0) grown on vertical agar plates in the presence of zinc (+Zn), in the absence of zinc (-Zn), or the absence of both phosphorus and zinc (-P-Zn).

### Regulation of cell cycle and cell proliferation may be an important biological process underlying -P-Fe’s compensation of -Fe mediated RGR reduction

To gain insight on the genetic architecture of the mechanisms that mitigate -Fe or - Zn’s inhibition of RGR by -P, we conducted GWAS to identify loci that were associated with the variation of RGR between double and single stresses. The traits we used are as follows: ∆RGR_(-P-Fe, -Fe)_ = RGR_(-P-Fe/Ct)_ - RGR_(-Fe/Ct)_, and ∆RGR_(-P-Zn, -Zn)_ = RGR_(-P-Zn/Ct)_ - RGR_(-Zn/Ct)_ (Figure 7 A-B). While the heritability for ∆RGR_(-P-Fe,-Fe)_ was estimated as 0.27, the ∆RGR_(-P-Zn,-Zn)_ trait missed heritability. Consistently, using a conservative Bonferroni threshold, 4 regions spanning 21 candidate genes were identified for ∆RGR_(-P-Fe,-Fe)_ (Supplemental Table 6), and no candidate genes were associated for ∆RGR_(-P-Zn,-Zn)_. From the 4 regions associated with ∆RGR_(-P-Fe,-Fe)_, 3 regions overlapped with QTLs identified in the previous analysis. The first region contains two genes, At2G05160 and AT2G05170, which were identified in the RGR for -P, -P-Fe and -P-Zn. The second and third regions span 7 and 8 genes, which were identified in the RGR for in -P-Fe and -P-Zn respectively. More importantly, the fourth region is specific to the ∆RGR_(-P-Fe,-Fe)_ trait, and contains 4 candidate genes including AT5G23980 that encodes a FERRIC REDUCTION OXIDASE 4, known to be involved in Fe reduction and absorption (Bernal et al., 2012).

**Figure 7.**
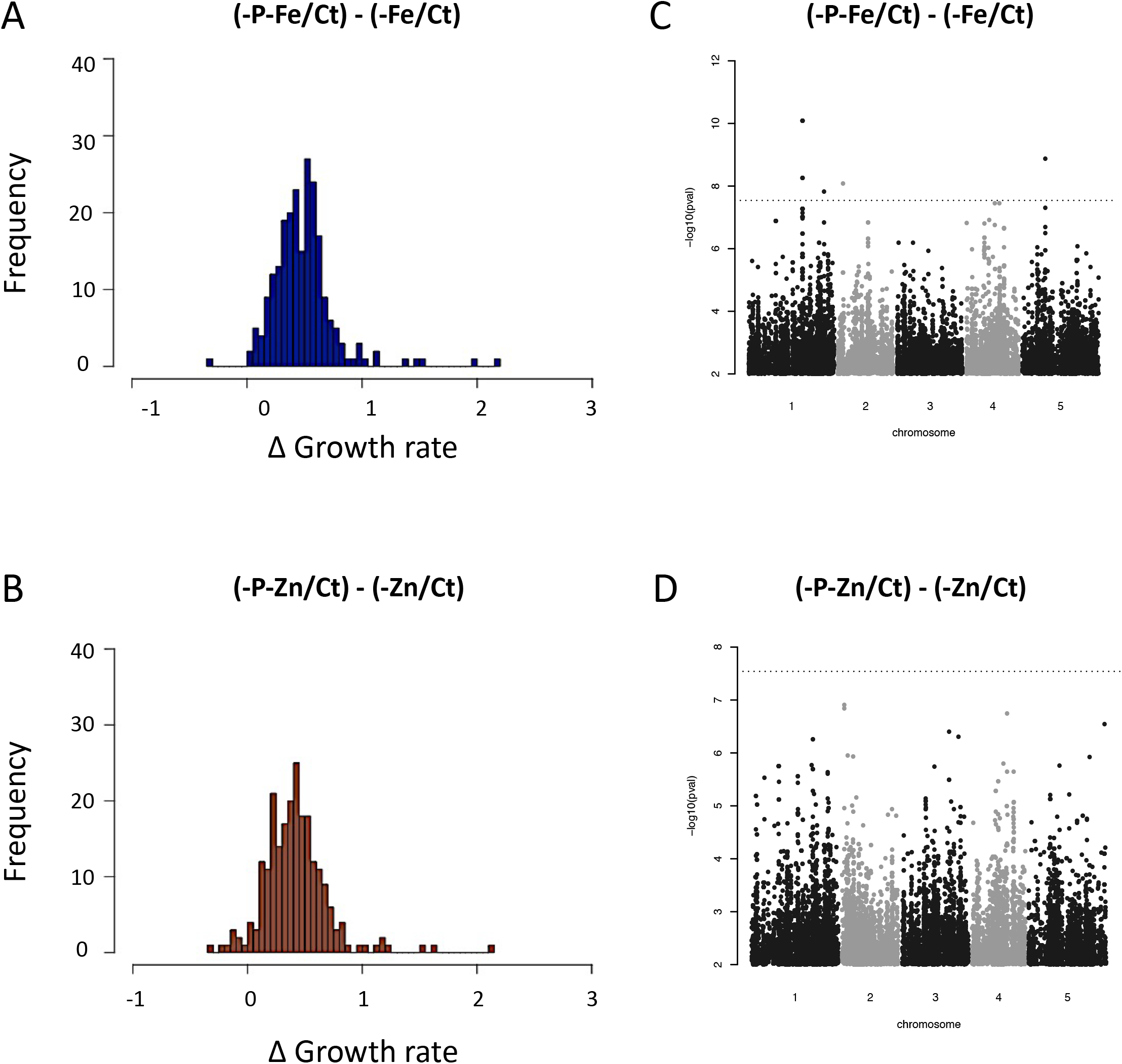
Compensation of root growth rate by nutrient limitation interactions and Manhattan plots of GWAS results. (A-B) Variation in the root growth rate (RGR) between -P-Fe and -Fe compared to control condition (Ct) presented as ∆RGR_(-P-Fe,-Fe)_= ((-P-Fe/Ct) - RGR(-Fe/Ct)), and (B) between -P-Zn and -Zn compared to control condition (Ct)-P-Zn and –Zn presented as ∆RGR_(-P-Zn,-Zn)_ = ((-P-Zn/Ct) - RGR(-Zn/Ct)). (C) SNPs associated with the ∆RGR_(-P-Fe,-Fe)_ or (D) ∆RGR_(-P-_ Zn, -Zn). SNPs are plotted according to their position along the appropriate chromosome. Plotting colors alternate between black and grey in order to facilitate the visualization of each of the five chromosomes. Bonferroni threshold is indicated by black dashed line. (C-D) Genome-wide distribution of the −log10 P-values of SNP/phenotype associations using the MMA method.

We further analyzed the genetic architectures of both traits, ∆RGR_(-P-Fe,-Fe)_ and ∆RGR_(-P-Zn,-Zn),_ by using a less stringent P value threshold. We chose −log10(P) < 4 threshold as it is frequently used in similar studies (e.g. Davila Olivas et al., 2016; Thoen et al., 2017). Our analysis found 949 and 437 associated markers (SNP) corresponding to 186 genes and 92 genes for ∆RGR_(-P-Fe,-Fe)_ and ∆RGR_(-P-Zn,-Zn)_ respectively (Supplemental Table 7). Among the genes detected for ∆RGR_(-P-Fe,-Fe)_, we found *VITAMINC4* (*VTC4*), a gene encoding a protein with dual myo-inositol-monophosphatase and ascorbate synthase activity involved in the reduction of Fe^+3^ to produce the Fe^+2^ (Torabinejad et al., 2009). Interestingly, VTC4 belongs to the set of genes that are direct targets of the PHOSPHARE RESPONSE1 (PHR1) (Bustos et al., 2010), and is considered as a potential candidate for the cross-talk between Fe and P signaling to regulate root growth (Mora-Macías et al., 2017). An overlap of 31 SNPs (Supplemental Table 8) that correspond to 10 genes was detected (Supplemental Table 9) was detected for ∆RGR_(-P-Fe,-Fe)_ and ∆RGR_(-P-Zn,-Zn)_. This overlap is significantly higher compared to random markers (p=2.8^-11^) or permutations for which no significant overlap at this threshold was observed. In this list we can distinguish in particular many genes involved in the transcriptional regulation (e.g. AT1G27730, *SALT TOLERANCE ZINC FINGER*; AT1G27660, *bHLH110*) and key gene involved in the DNA methylation (AT1G57820, *VARIANT IN METHYLATION 1*).

To capitalize upon our GWAS results, and to go beyond our *a priori* gene list to find pathways and functional modules that explain the variation of RGR, we took advantage of the existence of genome-scale gene co-function networks such as AraNetv2 that covers 84% of *A. thaliana*’s coding genes (Lee et al., 2015). Using this tool and all GWAS candidate genes for the ∆RGR(_-P-Fe_, _-Fe_) trait, we identified three modules (Supplemental Table 10). The largest module comprises 26 genes connected with one another (Figure 8). The Gene Ontology (GO) annotation analysis revealed that there is significant enrichment for “chromatin modification” (p=2.44*10- 6), “DNA replication” (p=9.58*10-5), and “regulation of cell cycle” (3.37*10-4) within this module. A similar analysis for genes associated with ∆RGR_(-P-Zn,-Zn)_ showed one module comprising 6 genes (Supplemental Table 12) with a significant GO enrichment for the “regulation of cell cycle” (p=2.8*10-6) and “cell proliferation” (p=3.85*10-6) (Supplemental Table 10). Taken together, our analyses suggest that similar molecular mechanisms may be involved in promoting RGR under different combined nutrient deficiency.

**Figure 8.**
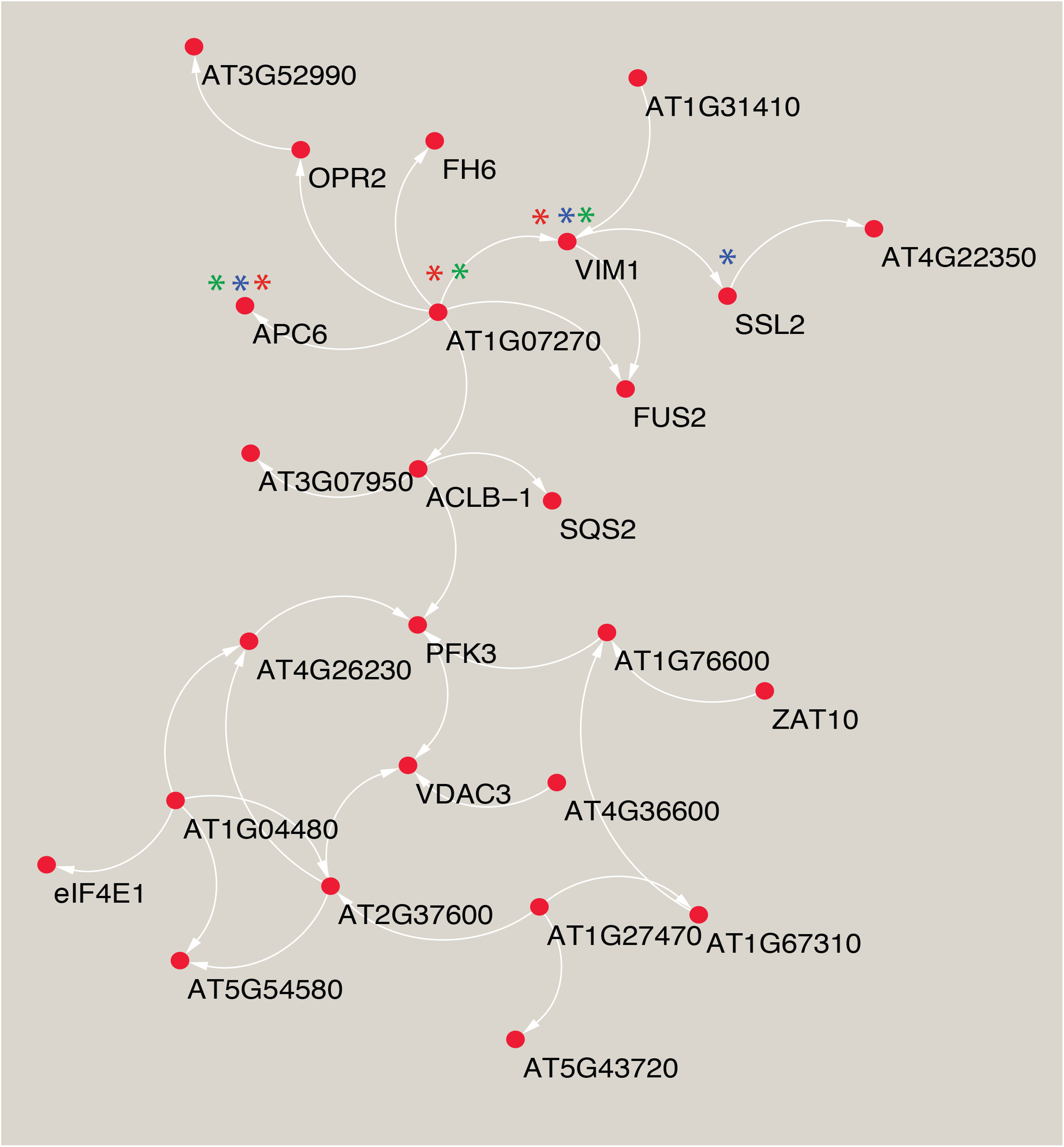
Predicted molecular pathway for the compensation of root growth rate by phosphate and iron combined stresses. A network between the subset of the GWAS candidate genes identified using the publicly available resource AraNet (Lee et al., 2015). The network visualization is generated using Cytoscape software (Shannon et al., 2003). The Gene Ontology (GO)-biological process enrichment of a subset list of GWAS genes for ∆RGR_(-P-Fe,-Fe)_ were conducted using AraNet. Genes enriched in GO terms are marked with stars: “chromatin modification” (blue), “DNA replication” (red), and “regulation of cell cycle ” (green).

## Discussion

Below ground, roots can sense and respond to the nutrient stress during all phases of plant growth and development (Schachtman and Shin, 2007). An important part of this response is adjustment of the root growth rate in response to nutrient availability (López-Bucio et al., 2003; Osmont et al., 2007; Kellermeier et al., 2014). This plasticity of root growth is an important adaptive trait. Nevertheless, despite its primary importance for optimizing root foraging for resources in heterogeneous soil environments and ensuring crop yield, most of the work on plant mineral nutrition has focused on responses to absence of single nutrients, and the few studies that went beyond this did examined these responses only in a single strain (e.g. Kellermeier et al., 2014). Here, we report the first extensive analysis of the RGR in 227 accessions of *A. thaliana* grown under six nutritional conditions including combinatorial nutrient deficiencies: control (MS, Ct), -P, -Fe, -Zn, -P-Fe, and -P-Zn. We focused on the variation of RGR in an early phase of plant development given its fundamental importance in the plant life cycle. We showed the presence of a large amount of heritable natural variation of RGR under each growth condition. We then performed 7 GWAS to look for the genetic variants underlying the observed natural variation of RGR, and present all significant associations. These data provide an insight into the genetic basis that underlies root growth responses to multiple nutrient deficiencies and lay a firm foundation to identify and characterize causative polymorphisms through functional molecular work.

Genetic differences between *A. thaliana* accessions underlie the plant’s extensive phenotypic variation, and until now these have been interpreted largely in the context of the Col-0 accession. While Col-0 has been the predominant natural accession for research in plant biology for many decades (Somssich et al., 2018). Our work provides clear evidence that Col-0 is not the best representative of the species for P and Zn deficiency responses, which is highly relevant because Col-0 responses were often assumed to be the general responses of plants to nutrient deficiencies. Unlike Col-0, most of the accessions reduce the RGR under -Zn, and more importantly most of the accessions stimulate the RGR under -P. This result indicates that Col-0 is not the ideal accession to study all types of nutrient stresses. In -P, while Col-0 consistently showed a reduction of primary root growth in many studies, the amplitude of such RGR reduction varied largely between studies. For example, in Reymond et al., (2006) Col-0 exhibited a strong response to -P (5 μM P), whereas in Chevalier et al. (2003) (1 μM P), the primary root is only slightly affected. Reversely, the Shadara accession showed invariably an oversensitivity to -P conditions (Reymond et al., 2006; Svistoonoff et al., 2007; Shahzad et al., 2018), which was also observed in our study. This response in Shadara was used to generate a RIL population (using Bay-0 displaying long root under -P compared to Shadara as the other parent) that allowed the mapping of a key gene regulating root growth under -P, *LPR1* (Reymond et al., 2006). Despite its contrasting and consistent root growth responses to -P compared to Col-0, Shadara remains little used to study responses to -P. In our study, we examined the RGR in the early stages of plant development in response to -P, and thereby revealed that many accessions sense and respond to depletion of P in the medium by stimulating RGR. As these responses were observed early on during plant development, when P levels in the seedlings will not be significantly depleted, our result suggests the existence of mechanisms that perceive changes in P and contribute to early RGR modulation by -P. These mechanisms remain to be discovered, but once discovered, might be exploited to design strategies to improve root growth under -P.

Most accessions displayed a reduction in the RGR under -Fe or -Zn conditions. However, the extent of this reduction varied between -Fe and -Zn with – Fe causing more reduction in general. Interestingly, when each of those single deficiency conditions was combined with -P, we observed an increase of RGR for most of the accessions. This indicates the existence of strong interdependencies of the root growth responses between the macronutrient P and micronutrients Fe and Zn. In line with our result, Rai et al., (2015) showed that the primary root length was significantly increased in response to -Fe-P compared to -Fe in Col-0. Furthermore, it has been well documented that -Fe or -Zn leads to the accumulation of P in plants (Rai et al., 2015; Khan et al., 2017; Kisko et al., 2018) likely through the activation of root Pi uptake (Huang et al., 2000,Kisko et al., 2018). Whether the accumulation of P in -Zn or -Fe conditions is directly related to the reduction of RGR will need further investigations. Conversely, plant over-accumulates Fe in -P conditions. As aforementioned, it has been proposed that the Fe over-accumulation in Col-0 roots could reach a “toxic” level that cause the inhibition of root growth in -P. Consistently, the primary root growth grown under simultaneous absence of Fe and P is comparable to those observed under control condition (Ward et al., 2008; Balzergue et al., 2017; Mora-Macias et al., 2017; Abel et al., 2017; Gutierrez-Alanis et al., 2018). Our results confirm the observation made for Col-0, and extend it to other accessions (e.g. Shadara, Sorbo, Rd-0, Rmx-A180, Mr-0, Hau-0, Got-22, Mdn-1, RRS-10). Nevertheless, interestingly, this rescue of RGR by the additional absence of -Fe was not observed in most of the accession that we tested. Many accessions exhibited a reduction of RGR by -P-Fe compared to -P alone. This result suggests that the availability of Fe in the medium is not the sole determinant that control RGR under -P condition. Identification of mechanism(s) that control root growth under simultaneous -P-Fe deficiency deserves further research.

Taken together, our results show that there are multiple, genetically determined strategies for how plants respond to nutrient limitations. It is therefore difficult to generalize observations made on one single accession such as Col-0 to the species level. A more comprehensive strategy to study a variety of accessions is currently becoming feasible thanks to the availability of the genome sequence of thousands of accessions, and the possibility to generate mutations in genes of interest via gene editing technologies. Our results further highlight the importance of studying combined nutrient stresses to comprehensively understand plant responses to nutrient deficiency stresses and their underlying genetic and molecular mechanisms.

Our data revealed an important aspect of plant responses to combined stresses. The overlap between GWAS candidate genes from two single stresses and their combination is very limited. This result strongly suggests that there is a distinct genetic architecture underlying the responses to single and combined nutrient stresses in plants. This is further supported by recent work exploring natural variation in Arabidopsis in response to abiotic (drought) or biotic (fungal pathogen) stresses, which indicated that distinct genetic mechanisms underlie responses to single and combined stresses (Davila Olivas et al., 2016). This finding is relevant for designing strategies to improve crop yields in the field, as natural environments impose multiple stresses at once. In particular, for gene stacking approaches deploying multiple genes or alleles to help crops growing in the field that face multiple stresses, our data are in favor of using multiple “specialized genes” for combined stresses rather than stacking genes addressing individual stresses (Halpin, 2005). The “specialized genes” could be revealed through quantitative genetics approaches such as GWAS.

Time and cost-efficiency of GWAS has made it a useful approach to understand the genetic and molecular factors that govern complex traits, such as the root growth under different nutritional conditions (Satbhai et al., 2017; Bouain et al., 2018). Our GWASs led to obtain lists of the most significant SNPs associated with the variation of RGR under each of the aforementioned six nutrient-deficient conditions. We then identified candidate genes covered by the GWAS QTLs. Some genes that are known to be involved in the regulation of Col-0 root growth under specific nutrient stress conditions were detected with modest p-values. For example, key genes known to regulate root growth under -P or -Fe have been detected, namely *HPS7 (p=3.83*10^-6^)* (Kang et al., 2014) for -P, and *bHLH104 (p=2.3*10^-6^)* for - Fe (Li et al., 2016). Interestingly, the well-characterized *LPR1* gene was identified among the GWAS candidate genes on RGR under -P (*p= 8.18*10^-5^*), -P-Fe (*p= 2.57*10^-5^*), and on ∆RGR (P-Fe, -Fe) (*p= 3. 2*10^-5^*). The *lpr1* mutant under –P is characterized by not only its longer primary root, but also its lower Fe content compared to wild-type plants (Balzergue et al., 2017). Nevertheless, whether the lower Fe content in *lpr1* is enough to trigger Fe deficiency signaling is unknown. If this hypothesis were confirmed, LPR1 would play an important role in integrating both Fe and P signals to control the primary root elongation. This hypothesis is supported by the recovery of root growth in Col-0 under combined -P and -Fe stresses compared to -P alone (Ward et al., 2008).

Beyond the root growth related genes, our GWAS identified many new candidate genes that were previously unknown to regulate RGR under single or combined nutrient deficiency. Nevertheless, it is important to go beyond the detection of candidate genes by GWAS either by functional validation of top ranked candidates based on p-values, or through to the identification of pathways and functional modules that explain complex traits. Starting with a list of candidate genes detected by GWAS for the compensation of RGR under -P-Fe/Ct compared to -Fe/Ct or under-P-Zn/Ct compared to -Zn/Ct described in this work, and by using a genome-scale gene co-function network AraNet (Lee et al., 2014), we predicted pathways that might underlie the compensation strategy. Four statistically enriched biological processes included the regulation of cell cycle, chromatin modification, cell proliferation, and DNA replication. These biological processes can be subjected to perturbation to validate their role in the compensatory mechanism seen under -P-Fe or -P-Zn for RGR. Such systems genetics strategy will accelerate future research discovery leading to the improvement of our understanding of the regulation of root growth under single and combined nutrient deficiency.

## Materials and Methods

### Plant materials and growth conditions

A total of 227 natural accessions of *A. thaliana* (S Table 1) were phenotyped and used to perform GWAS. Six growth conditions were used, MS (Control; Ct), phosphate deficiency (-Pi), iron deficiency (-Fe), zinc deficiency (-Zn), phosphate and iron deficiency (-Pi-Fe) and phosphate and zinc deficiency (-Pi-Zn). Seeds were surface-sterilized in chlorine gas for 1□h. Chlorine gas was generated from 130□ml of 10% sodium hypochlorite and 3.5□ml of 37% hydrochloric acid. After sterilization, seeds were imbibed in water and stratified for 3 days at 4□°C in the dark to promote uniform germination. For each genotype we sowed 12 seeds, distributed on 4 different plates. Each plate contained eight different accessions with 3 plants per accession. Plates were placed in the growth chamber in a randomized manner. Seeds were then germinated and grown vertically on 1X MS-agar medium, which contained 1 mM KH_2_PO_4_, 1 mM MgSO_4_, 0.5 mM KNO_3_, 0.25 mM Ca(NO_3_)_2_, 10 μM MnCl_2_, 30 μM H_3_BO_3_, 1 μM CuCl_2_, 0.1 μM (NH_4_)_6_Mo_7_O_24_, 50μM KCl, 100 μM NaFeEDTA and 15 μM ZnSO_4_ in presence of 0.8% (wt/vol) agar and 1 % (wt/vol) sucrose. -Pi medium was made by replacing the source of Pi (KH_2_PO_4_) to with 1 mM KH_2_CaCO_3_. -Zn medium was made by omitting the source of Zn (ZnSO_4_) in the medium. -P-Zn medium was made by not adding the only source of Zn (ZnSO_4_) and by replacing KH_2_PO_4_ with 1 mM KH_2_CaCO_3_ to the medium. -Fe medium was made by not adding the FeEDTA in the medium and by supplying the FerroZine that is known as a strong Fe chelator. -P-Fe medium was made by not adding the source of Fe (FeEDTA) and by supplying the FerroZine, and by replacing P source (KH_2_PO_4_) by 1mM KH_2_CaCO_3_ to the medium. Plants were grown at 22°C in the same growth chambers under the same light regime (long-day: 8 h dark, 16 h light) conditions. Plant phenotyping for GWAS was performed as described previously (Slovak et al., 2014). The BRAT software was used to perform root trait quantification (Slovak et al., 2014).

### Genome wide association studies (GWAS)

Genome-wide association mapping was performed on the regression coefficients of root growth of three-, four- and five-days-old seedlings grown under the above detailed conditions. The phenotypic data for the rare root growth and the regression coefficients used in the analysis are available at the AraPheno database (Seren et al., 2016). The genotypic data were based on whole genome sequencing data [The 1001 Genomes Consortium, 2016] and covered 4,932,457 SNPs for the 227 accessions. 1,739,142 of these markers had a minor Allele frequency of at least 5% in the population and where further used for GWAS. GWAS was performed with a mixed model correcting for population structure in a two-step procedure, where first all markers where analyzed with a fast approximation (emmaX, [Kang et al. 2010]) and afterwards the top 1000 markers where reanalyzed with the correct full model. The kinship structure has been calculated under the assumption of the infinitesimal model using all genetic markers with a minor Allele Frequency of more than 5 % in the whole population. The analysis was performed in R (R Core Team (2016)). The used R scripts are available at https://github.com/arthurkorte/GWAS. The Genotype Data used for GWAS are available www.1001genomes.org.

Heritability estimates have been extracted from the mixed model according to the formula: H^2^ = V_G_ /(V_G_ + V_E_), where V_G_ is the among-genotype variance component and V_E_ is the residual (error) variance.

### Molecular pathway prediction

The functional modules were predicted based on the GWAS genes using the publicly available resource AraNet (Lee et al., 2015). Network visualization was generated using Cytoscape software (Shannon et al., 2003).

## Acknowledgment

The authors are thankful to Bonnie Wohlrab and Christian Göschl for technical assistance. This work was funded by the “Institut National de la Recherche Agronomique – Montpellier - France” INRA and by the AgreeenSkills Plus to H.R., and supported by funds from the Austrian Academy of Science through the Gregor Mendel Institute (GMI) to W.B., an Austrian Science Fund (FWF) stand-alone project (P27163-B22) to W.B., and by the National Institute of General Medical Sciences of the National Institutes of Health (grant number R01GM127759 to W.B.), and supported by funds from the Carnegie Institution for Science and Brigitte Berthelemot to S.Y.R.

## Author Contributions

H.R. coordinated the project and writing of the manuscript. H.R. and W.B. designed the GWAS. N.B. and S.B.S. carried out the experiments. A.K. performed the GWAS analysis. S.Y.R. and H.R. conceived data analysis scheme, analyzed the data, and formulated the manuscript.

## Declaration of interest

The authors declare no competing financial interests.

---

**Supplemental Table 1.** Primary root length and root growth rate (RGR) of 227 natural accessions of *Arabidopsis thaliana* grown on control condition (Ct), deficiency of phosphorus (-P), iron (-Fe), zinc (-Zn), phosphorus and iron (-P-Fe), or phosphorus and zinc (-P-Zn). Measurements were done on 3, 4 and 5 days-old seedlings.

**Supplemental Table 2.** Normalized root growth rate (RGR) of the 227 natural *Arabidopsis thaliana* accessions. Values were obtained by diving the RGR presented in Supplemental Table 2 for each of the nutrient-deficient conditions with the RGR under control condition (Ct).

**Supplemental Table 3.** Heritability estimates for the root growth rate (RGR) and the normalized RGR using estimates from the linear mixed model.

**Supplemental Table 4.** List of 145 candidate genes that are present in a 10kb window around significant SNPs for the GWAS of normalized root growth rate (RGR) under nutrient-deficient conditions. The threshold used to declare these SNPs significant is Bonferroni.

**Supplemental Table 5.** List of 87 significant SNPs for the GWAS of normalized root growth rate (RGR) under the different nutrient-deficient conditions and the genes that are surrounding these SNPs

**Supplemental Table 6.** List of 21 candidate genes found in the analyses of root growth rate (RGR) of phosphorus and iron deficiency (-P-Fe) normalized on iron deficiency (-Fe).

**Supplemental Table 7.** List of candidate genes found in the GWAS analyses (threshold of p=10^-4^) of root growth rate (RGR) of phosphorus and iron deficiency (-P-Fe) normalized on iron deficiency (-Fe) and phosphorus and zinc deficiency (-P-Zn) normalized on zinc deficiency (-Zn) .

**Supplemental Table 8.** List of 31 SNPs that are found in the GWAS analyses (threshold of p=10^-4^) of RGR of phosphorus and iron deficiency (-P-Fe) normalized on iron deficiency (-Fe) and phosphorus and zinc deficiency (-P-Zn) normalized on zinc deficiency (-Zn).

**Supplemental Table 9.** List of 10 genes that are found in both, the analyses of RGR of phosphorus and iron deficiency (-P-Fe) normalized on iron deficiency (-Fe) and phosphorus and zinc deficiency (-P-Zn) normalized on zinc deficiency (-Zn) at a less stringent threshold of p=10^-4^.

**Supplemental Table 10.** List of genes forming the modules identified through the analyses of GWAS candidate genes for ∆RGR_(-P-Fe,-Fe)_ and for ∆RGR_(-P-Zn,-Zn)_ traits using the publicly available resources, AraNet.

